# A rare natural lipid induces neuroglobin expression to prevent amyloid oligomers toxicity and retinal neurodegeneration

**DOI:** 10.1101/2021.06.23.449608

**Authors:** Henry Patrick Oamen, Nathaly Romero Romero, Philip Knuckles, Juha Saarikangas, Yuhong Dong, Fabrice Caudron

## Abstract

Most neurodegenerative diseases such as Alzheimer’s disease are proteinopathies linked to the toxicity of amyloid oligomers. Treatments to delay or cure these diseases are lacking. Using budding yeast, we report that the natural lipid tripentadecanoin induces expression of the nitric oxide oxidoreductase Yhb1 to prevent the formation of protein aggregates during aging and extends replicative lifespan. In mammals, tripentadecanoin induces expression of the Yhb1 orthologue, neuroglobin, to protect neurons against amyloid toxicity. Tripentadecanoin also rescues photoreceptors in a mouse model of retinal degeneration and retinal ganglion cells in a Rhesus monkey model of optic atrophy. Together, we propose that tripentadecanoin affects p-bodies to induce neuroglobin expression and offers a potential treatment for proteinopathies and retinal neurodegeneration.

**One Sentence Summary:** The natural lipid tripentadecanoin is cytoprotective against amyloid oligomer toxicity and retinal neurodegeneration by inducing YHBI/neuroglobin expression in yeast and mammals.

## Main text

Age-associated neurodegenerative diseases, including Alzheimer’s, Parkinson’s and prion diseases are linked to the toxicity caused by protein misfolding, particularly into amyloid fold (*1*). For example, brains of Alzheimer’s disease patients commonly display β-amyloid and/or neurofibrillary tangles of tau that accumulate outside and inside neurons, spread between cells and thereby disrupt normal cell functions. These diseases are becoming increasingly prevalent and current curative therapies are insufficient. Therefore, there is an urgent need to develop therapeutic agents that counteract amyloid toxicity.

*Ophioglossum thermale* is a medicinal herb used in traditional Chinese medicine as a rescuing treatment for snake bite toxicity and occasionally for conditions of neuronal atrophy. The glycerolipid tripentadecanoin (Figure 1A) was reported as one of *Ophioglossum’s* extract components (*2*). We tested the effect of *Ophioglossum* whole extracts and tripentadecanoin on neurons challenged by β-amyloid_1-42_ oligomers (AβO) toxicity. Mouse primary cortex neurons were incubated 48 hours with different concentrations of the herb whole extract, tripentadecanoin or docosahexaenoic acid (DHA, 22:6 n-3) as a positive control (*3*) before addition of 1μM AβO. After 24 hours, cell viability was assayed by MTT-colorimetry and demonstrated that the herb extract provided neuroprotection against AβO at all concentrations tested (Figure 1B). Tripentadecanoin provided neuroprotection from 100nM and higher concentrations (Figure 1C), indicating that it is the active compound that conferred the neuroprotective effect of *Ophioglossum* extract. Testing other lipids with similar structures using this assay demonstrated that the neuroprotection effect was specific to tripentadecanoin, although linoleic acid provided some neuroprotection (Supplemental Figure 1). We then asked whether tripentadecanoin was also neuroprotective when added concurrently or after AβO. For these experiments, we used Humanin (HNG) as a positive control (*4*), since docosahexaenoic acid does not protect neurons in these conditions. Tripentadecanoin was added at 0, 3 or 6 hours after AβO. Remarkably, tripentadecanoin was highly neurorescuing (320nM and 1μM) after 3 hours and neurorescuing after 6 hours (Figure 1D). Next, we tested tripentadecanoin neuroprotective effect on human cells. We used neurons derived from induced pluripotent stem cells and challenged them with AβO (1μM). Tripentadecanoin or HNG was added after 0, 3 or 6 hours and cell viability was assayed by a neuron specific enolase assay. Compared to mouse neurons, tripentadecanoin protected human neurons with a higher efficiency (Figure 1E). We conclude that *Ophioglossum* extract and tripentadecanoin are neuroprotective and neurorescuing pharmaceutical agents against AβO induced toxicity. To test if the protective effect was specific to AβO or more broadly applicable to toxic amyloids, we tested its effects in mouse primary cortex neurons challenged with human α-synuclein oligomers (1μM), human amylin oligomers (1μM), Prion Protein_118-135_ oligomers (2μM) and human Tau oligomers (1μM). Importantly, tripentadecanoin displayed neuroprotective effects for all these toxic proteins (Figure 1F) suggesting a common downstream target.

**Figure 1.**
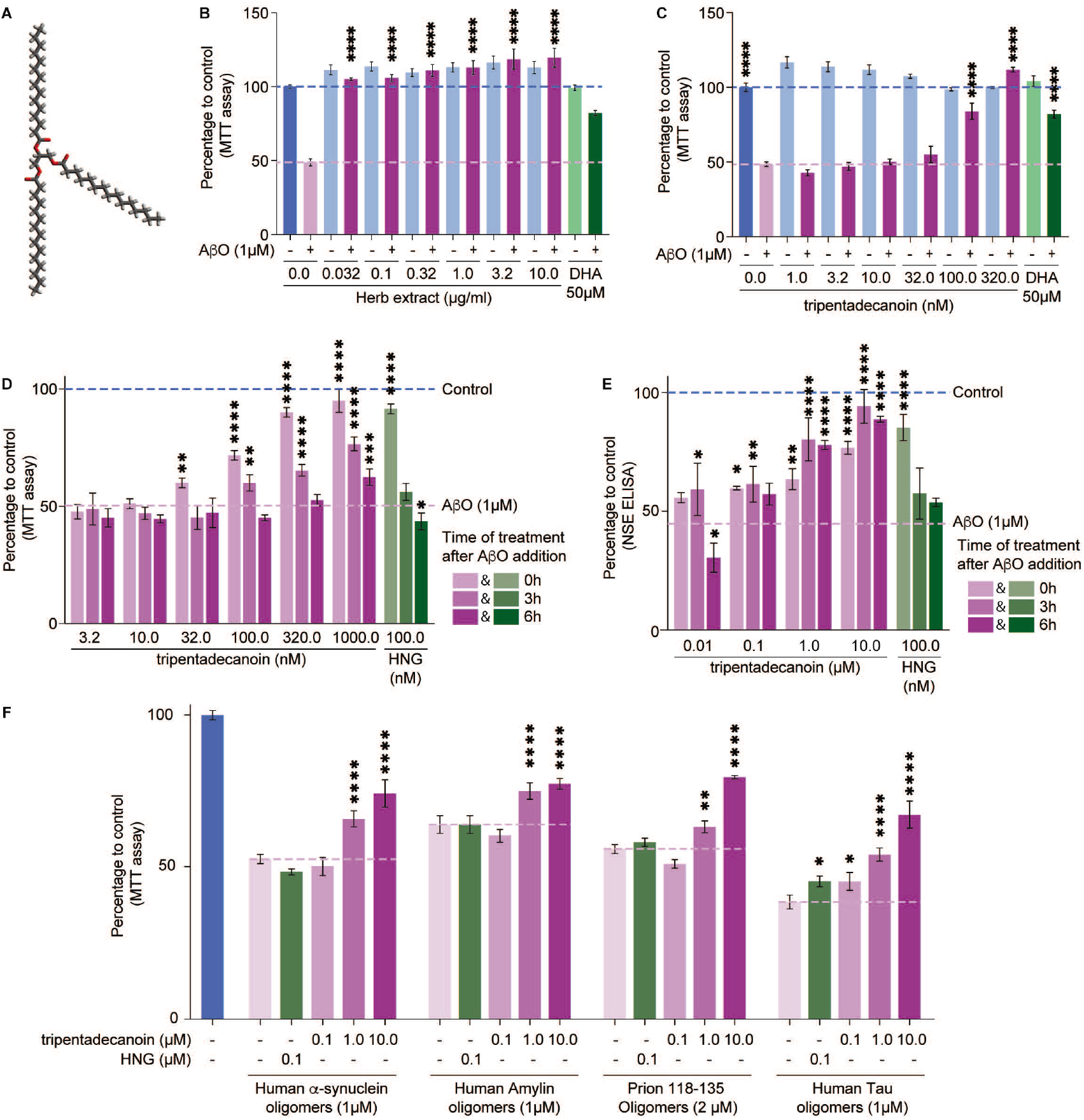
Ophioglossum whole extract and tripentadecanoin protect neurons against amyloid toxicity. **A.** Structure of tripentadecanoin. **B-C.** Quantification of mouse primary cortex neurons viability (MTT assay) after exposure to AβO (1μM) and pre-treatment with Ophioglossum extract or DHA (**B**) or pre-treatment with tripentadecanoin or DHA (**C**). **D**. Quantification of mouse primary cortex neurons viability (MTT assay) after treatment with tripentadecanoin or HNG 0h, 3h or 6h after exposure to AβO (1μM). **E**. Quantification of human induced pluripotent stem cells derived neurons viability (NSE assay) after treatment with tripentadecanoin or HNG 0h, 3h or 6h after exposure to AβO (1μM). **F**. Quantification of mouse primary cortex neurons viability (MTT assay) after treatment with tripentadecanoin or HNG 3h after exposure to human α-synuclein oligomers (1μM), human amylin oligomers (1μM), Prion Protein_118-135_ oligomers (2μM) or human Tau oligomers (1μM). P-values were obtained from ANOVA comparing to the toxin only treatment (A-E); *<0.05; **<0.01; ***<0.001; ****<0.0001.

To understand the mode of action of tripentadecanoin, we used *Saccharomyces cerevisiae* as a model organism. Age-induced protein deposits are formed during yeast replicative ageing and can be detected in live cells expressing the protein disaggregase Hsp104 fused to a green fluorescent protein (GFP) as a single focus in the cytoplasm (*5, 6*). Age-induced protein deposits accumulate damaged proteins and limit replicative lifespan (*5, 7, 8*). We thus used the age-induced protein deposit as a readout for activity of tripentadecanoin counteracting the formation and toxicity of damaged and misfolded proteins. We obtained 10 generations old cells expressing Hsp104-GFP from its endogenous locus, cultured in liquid media. Remarkably, old cells exposed to 1μM, 10μM and 30μM tripentadecanoin throughout the ageing process were less prone to display a Hsp104-GFP labelled protein aggregate, in a dose dependent manner, than untreated cells of the same age (Figure 2A-B). We observed a similar result with a whole extract from *Ophioglossum* (10μg/ml). Interestingly, exposure of young cells to tripentadecanoin (30μM) for 3 hours at the beginning of the experiment led to a similar inhibition of protein aggregate formation as when cells are exposed throughout their ageing (Figure 2C). In contrast, exposure of old cells to tripentadecanoin (30μM) for 3 hours just before imaging did not reduce the proportion of cells with Hsp104-GFP foci (Figure 2C). Remarkably, prevention of the formation of age-induced protein aggregate was accompanied by an increase in replicative lifespan (Figure 2D).

**Figure 2.**
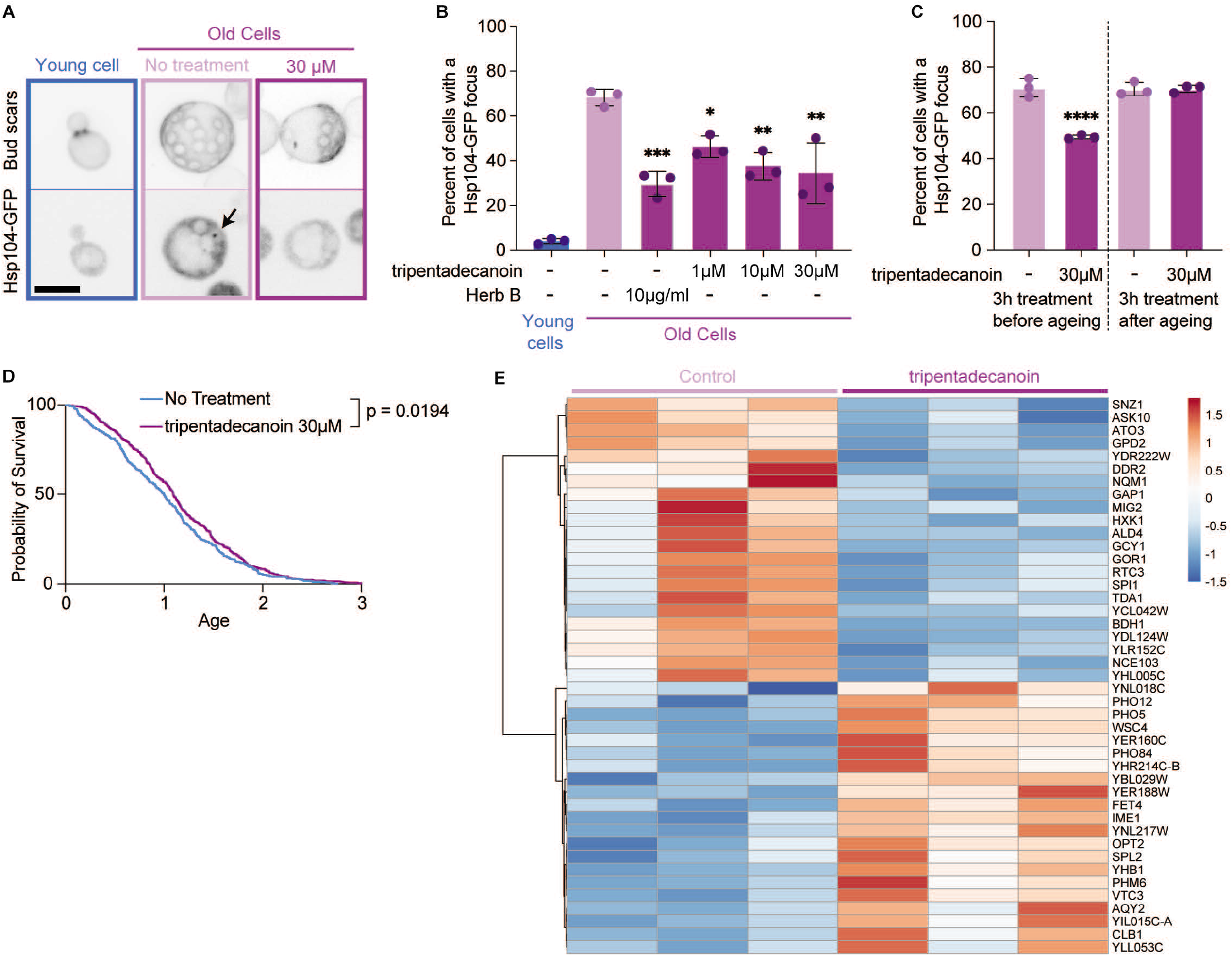
Tripentadecanoin prevents the formation of age-induced protein deposits in budding yeast. **A**. Representative images of a young cell, or old cells ± tripentadecanoin (30 μM). Upper panels show cells stained with fluorescent brightener 28 to reveal bud scars. Lower panels display the Hsp104-GFP signal. The arrow points at an age-induced protein deposit. Scale bar = 5μm. **B**. Percentage of cells with an Hsp104-GFP focus. Mean ±SD. Dots represent independent experiments (n ≥ 75 cells). P values are adjusted p values from an ANOVA comparing to untreated old cells *<0.05; **<0.01; ****<0.0001. **C**. Percentage of cells exposed for 3 hours to tripentadecanoin before or after ageing with an Hsp104-GFP focus. Mean ±SD. Dots represent independent experiments (n ≥ 124 cells). P values are adjusted p values from an ANOVA comparing to untreated old cells ****<0.0001. **D**. Replicative lifespan analysis of yeast cells ±tripentadecanoin (30μM). Age is expressed as the area of microcolonies normalised to the median of untreated cells. n ≥ 379 microcolonies. P-value was obtained from a Log-rank (Mantel-Cox) test. **E**. Heat map of the most significantly differentially expressed genes ± tripentadecanoin (30μM).

The formation of these naturally occurring protein deposits is thus prevented by tripentadecanoin and allowed us to probe the mode of action of this compound. To identify genes that are differentially expressed in the presence of Tripentadecanoin and thereby possibly confer its cytoprotective and lifespan extending effects, we performed an RNAseq analysis (Figure 2E). Gene ontology analysis revealed that cellular response to oxidative stress, transmembrane transport, water transport, glycerol metabolic process and polyphosphate metabolic process were the most significantly enriched in the differentially expressed genes (Supplemental table 1). Notably, this included *YHB1,* the yeast orthologue of human neuroglobin, several genes related to phosphate metabolism (*PHO5, PHO84, PHO12, PHO8, PHO81* and *PHO86*), inorganic polyphosphate synthesis and transport (*VTC1, VTC2, VTC3* and *VTC4).* To further test the involvement of these genes in preventing the formation of age-induced protein deposits, we used knock-out strains of selected genes that included *YHB1, VTC4* and *PHO84.* Compared to wild type old cells, more *yhb1*Δ and *vtc4*Δ old cells contained an Hsp104-GFP focus, while *pho84*Δ old cells were similar to wild type old cells (Figure 3A-B). In addition, *yhb1*Δ and *vtc4*Δ cells typically contained multiple Hsp104-GFP foci (Figure 3A). These results suggest that both Yhb1 and Vtc4 counteract the formation of age induced protein deposits. We next tested whether tripentadecanoin was still preventing the formation of age-induced protein deposit in the mutant strains. While tripentadecanoin (30μM) reduced the percentage of old wild type and *pho84*Δ cells with a Hsp104-GFP focus, this effect was lost in *yhb1*Δ and *vtc4*Δ cells (Figure 3A-B). Thus, we conclude that both Yhb1 and Vtc4 are essential for the protective effect of tripentadecanoin. We further focused on Yhb1 because its mammalian orthologue, neuroglobin, is identified, while Vtc4’s orthologue is not.

**Figure 3.**
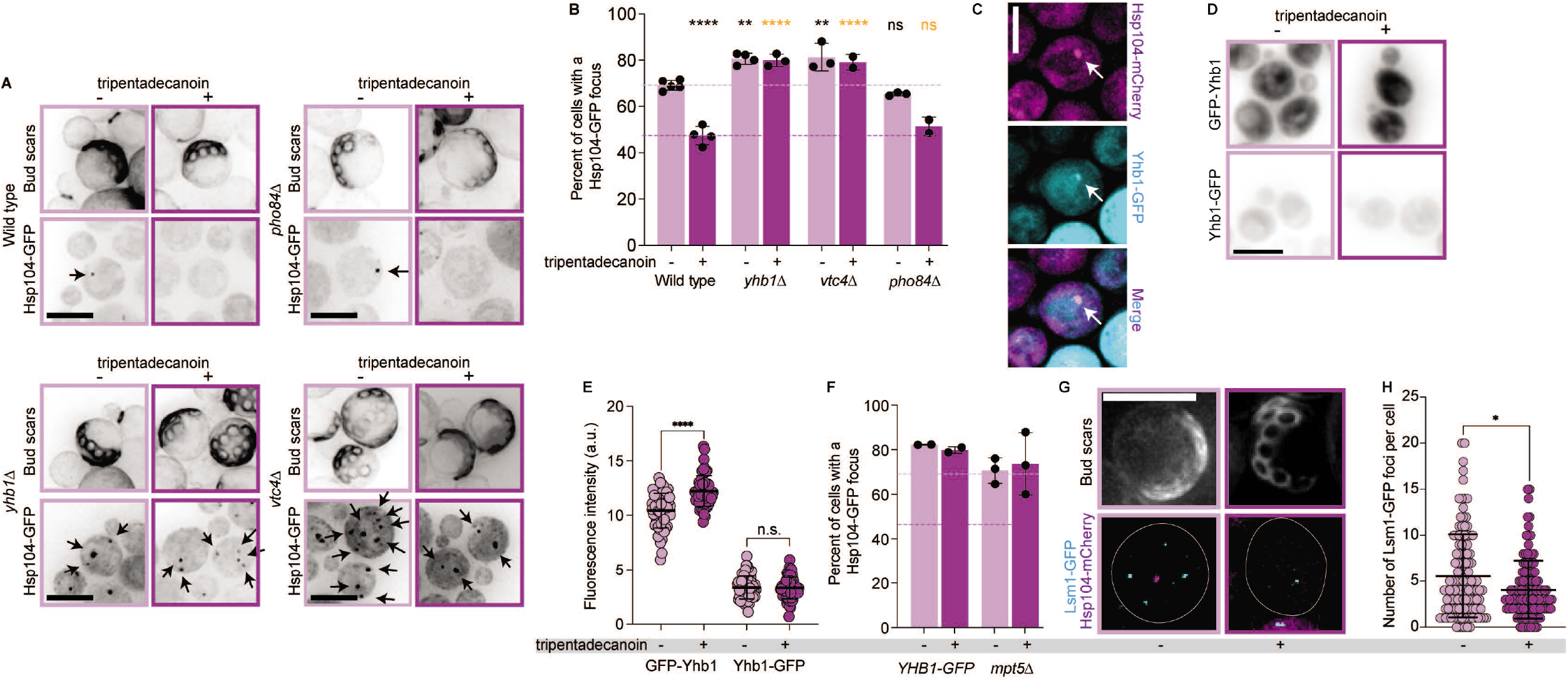
YHB1 mRNA regulation by P-bodies is required for tripentadecanoin to prevent the formation of age-induced protein deposits. **A**. Representative images of old cells ±tripentadecanoin (30μM). Bud scars stained with fluorescent brightener 28 (upper panels). Hsp104-GFP fluorescence signal (lower panels). Arrow point at age-induced protein deposits. Scale bars = 5μm. **B**. Percentage of cells with an Hsp104-GFP focus in the indicated genotypes. Mean ±SD. Dots represent independent experiments (n ≥ 211 cells). P values are adjusted p values from an ANOVA comparing to wild type control (black stars) or to wild type treated with tripentadecanoin (orange stars). **<0.01; ****<0.0001; ns = not significantly different. **C**. Colocalisation of Yhb1-GFP and Hsp104-mCherry foci in old cells. Scale bar = 5μm. **D**. Representative sum projection images of cells expressing Yhb1-GFP or GFP-Yhb1 ±tripentadecanoin. Scale bar = 5μm. **E**. Quantification of GFP-Yhb1 and Yhb1-GFP mean fluorescence intensity ± 30μM tripentadecanoin for 5 hours. Mean ±SD, n = 60 cells. p-values obtained from an unpaired t-test. ****<0.0001; ns = not significantly different. **F**. Percentage of cells with an Hsp104-GFP focus. Mean ±SD. Dots represent independent experiments (n ≥ 188 cells). Dotted lines correspond to the WT ±tripentadecanoin from Figure 3B. **G**. Representative images of old cells with bud scars stained with fluorescent brightener 28 (upper panels) and Lsm1-GFP and Hsp104-mCherry signal (lower panels). Scale bar = 5μm. **H**. Quantification of the number of Lsm1-GFP foci in old cells from Figure 3G. p-value = 0.0277 (*) obtained from a Mann-Whitney test (n > 123 cells).

We next tested whether Yhb1 is recruited to the age-induced protein deposit by obtaining old cells that express Yhb1-GFP and Hsp104-mCherry from their endogenous loci. 69.1% of the Hsp104-mCherry foci also recruited Yhb1-GFP (Figure 3C). This recruitment was not affected by treatment with 30μM tripentadecanoin (66.7% of the Hsp104-mCherry foci recruited Yhb1-GFP). We then asked if and how tripentadecanoin could induce *YHB1* expression. *YHB1* mRNA is destabilized by the Pumilio family proteins (Puf), particularly Mpt5 (Puf5) (*9*). This regulation occurs through two overlapping Puf recognition elements in the 3’untranslated region of *YHB1* mRNA. To test if the 3’UTR of *YHB1* was required for its induction by tripentadecanoin, we used the C-terminally tagged version of *YHB1* which replaces the native 3’UTR with the *ADH1* terminator (Yhb1-GFP) and a strain expressing a N-terminally tagged *YHB1* under the control of the strong *GPD* promoter (*10*) with the native *YHB1* terminator (GFP-Yhb1). After 5 hours of exposure to tripentadecanoin (30μM) expression of GFP-Yhb1 was induced but not of Yhb1-GFP (Figure 3D-E). Indeed, tripentadecanoin did not reduce the percentage of cells with an age-induced protein deposit in the Yhb1-GFP strain (Figure 3F). To test how tripentadecanoin induces *YHB1* expression further, we knocked out *MPT5* and obtained old cells expressing Hsp104-GFP. While *mpt5*Δ old cells harboured as many age-induced protein deposits as old wild type cells, tripentadecanoin had no effect on these cells (Figure 3F). These results suggested that induction of *YHB1* expression occurs at the level of its mRNA processing and prompted us to analyse the localisation of p-bodies in old cells with or without tripentadecanoin treatment. We tagged the p-body component Lsm1 with GFP and observed bright foci as described (*11*). However, after tripentadecanoin treatment, less Lsm1-GFP p-bodies were assembled in old cells (Figure 3G-H), suggesting that tripentadecanoin affects p-bodies regulation of *YHB1* mRNA decay/availability for translation in budding yeast.

To test whether tripentadecanoin also induces neuroglobin in mammalian cells, we measured the mRNA levels of neuroglobin in mouse primary cortex neuron. After 3 hours of treatment, neuroglobin mRNA level reached to 1.34 folds of the control for a 100nM treatment and 5.91 folds for a 1μM treatment (Figure 4A). These results strongly suggests that mRNA levels of neuroglobin are regulated similarly to *YHB1* in response to tripentadecanoin. To test the effect of tripentadecanoin in vivo, we chose an N-nitroso-N-methylurea (NMU) induced photoreceptor degeneration model for several reasons: Yhb1 has mostly been associated with nitrosated stress (*12*) while over-expression of neuroglobin was previously shown to rescue visual defects induced by NMU (*13*). Moreover, NMU causes photoreceptor degeneration within 7 days of a single-dose injection and the induced damage mimics the photoreceptor degenerative process in progressive human retinal degenerative diseases. Mice treated with NMU and tripentadecanoin (20 mg/kg or 50 mg/kg) displayed a significant protection of the outer nuclear layer of the retina compared to mice treated only with NMU (Figure 4B-D) well in agreement with the above mode of action studies. Furthermore, Harlequin mutant mice that display retinal ganglion cell loss and optic atrophy, due to a respiratory chain complex I defect, have a 2-fold reduced neuroglobin expression. Neuroglobin overexpression in harlequin mice eyes rescued retinal ganglion cell (RGC) body number and RGC axon number (*14*). Interestingly, following tripentadecanoin treatment in two rhesus monkeys (*Macata mulatta)* with unilateral optic atrophy (from a family with optical atrophy and retinal vascular abnormalities history, see material and methods), an increase in the average thickness of Retinal Nerve Fiber Layer (RNFL) was observed in the eyes with optic atrophy, but not in the healthy eyes (Figure 4E).

**Figure 4:**
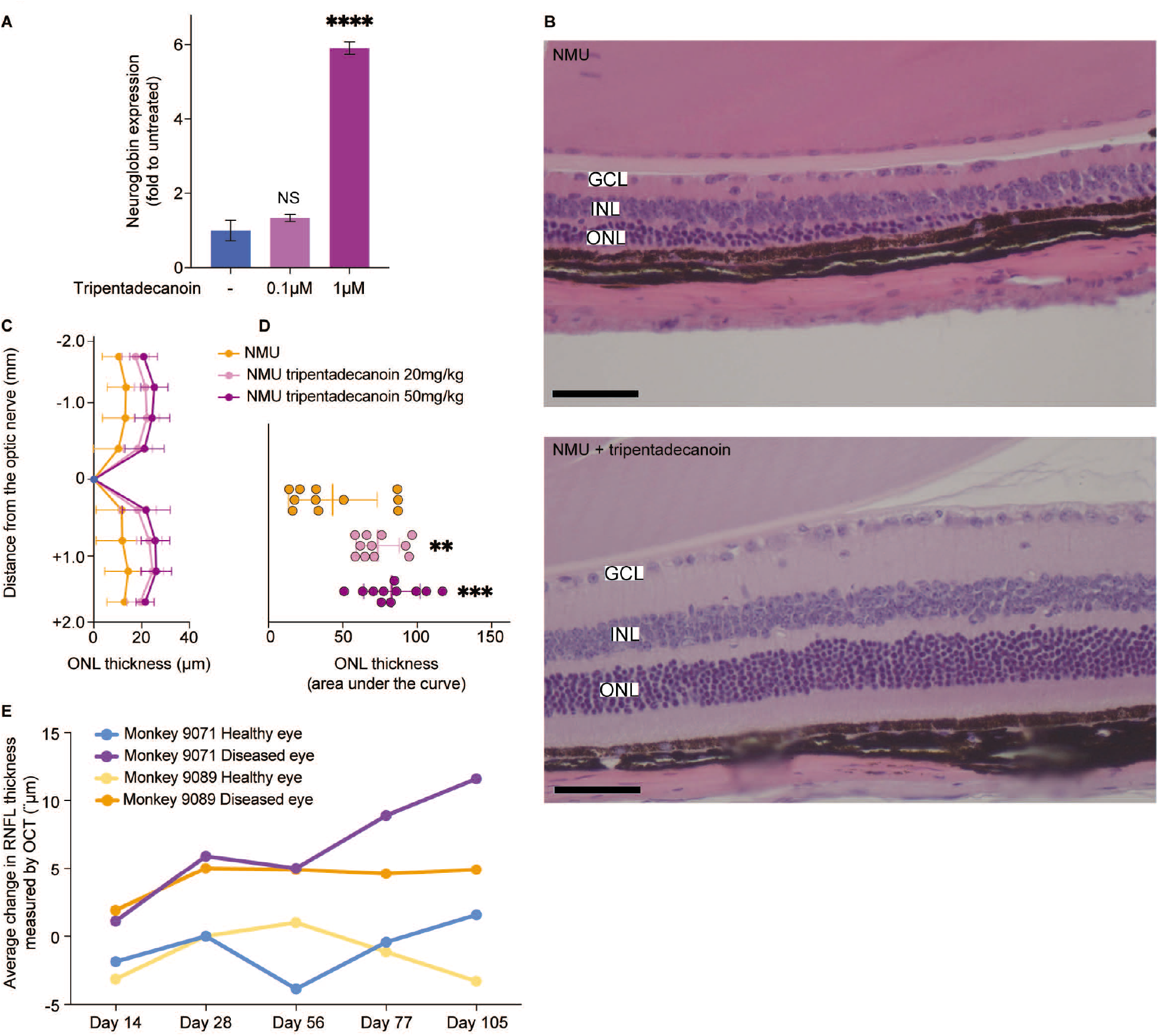
Tripentadecanoin induces neuroglobin expression in mouse primary cortex neurons and rescues NMU-induced photoreceptor damage in mice. **A.** Expression of neuroglobin mRNA over the control Rps28 mRNA presented as fold increase over untreated cells. P values are adjusted p values from an ANOVA comparing to untreated cells. ****<0.0001 **B.** Hematoxylin- and eosin-stained retinal sections of eyes from mice treated with NMU (top) or NMU + Tripentadecanoin (bottom). ONL: Outer nuclear layer, INL: Inner nuclear layer, GCL: Ganglion cell layer. Scale bar = 50μM **C.** Outer nuclear layer thickness measured from the retina sections as a function of the distance to the optical nerve (mm). Mean ±SD are presented. **D.** Outer nuclear layer thickness presented as the area under the curve from panel C. P values are adjusted p values from an ANOVA comparing to NMU treated cells. **<0.005; ***<0.001; ****<0.0001. **E**. Average change of retinal nerve fibre layer measured by OCT in monkeys #9071 and #9089 treated with tripentadecanoin. In each case, the left healthy eye and right diseased eye were measured.

Our results provide evidence that tripentadecanoin induces *YHB1* in budding yeast which prevents the formation of age induced protein aggregates. In mammals, we found that tripentadecanoin induces the Yhb1 orthologue neuroglobin, protects or rescues cells against toxic amyloids and prevents NMU-induced photoreceptor damage in mice and optic atrophy in Rhesus monkeys. Neuroglobin induction was previously shown to be protective during hypoxic–ischemic insults (*15*), to have cytoprotective effects against α-synuclein (*16*) and amyloid-β toxicity (*17*) and to inhibit apoptosis (*18*). Accordingly, its levels are increased in Alzheimer patients at early/moderate stages of the disease but decreased in severe cases (*19*). Neuroglobin appears as a central target to prevent different forms of neuronal degeneration. With tripentadecanoin, we have identified a small molecule that may protect patients against neurodegeneration of the retina as well as in proteinopathies.

---

**Supplemental Figure 1:**
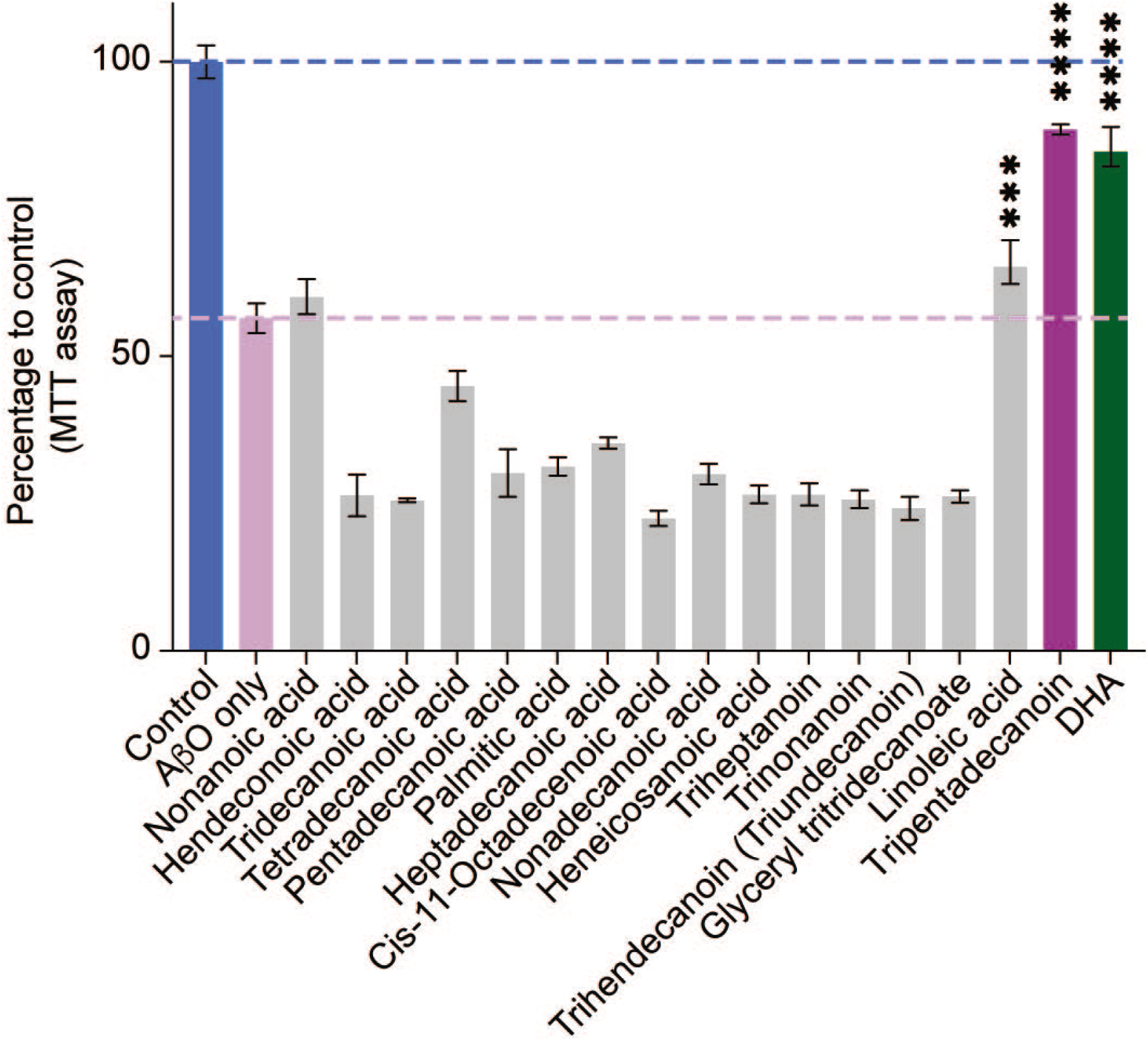
Quantification of mouse primary cortex neurons viability (MTT assay) after exposure to AβO (1μM) and pre-treatment with tripentadecanoin (1μM), DHA (50μM) or the indicated compounds at 1μM. P-values were obtained from ANOVA comparing to the toxin only treatment; ***<0.001; ****<0.0001.

**Supplemental Table 1:** List of Gene Ontology enrichments from the yeast RNAseq data.

**Supplemental Table 2:** yeast strain list.

**Supplemental Table 3:** RNAseq individual mRNA counts.

## Materials and methods

### Preparation of *Ophioglossum* whole extract

1 ml of DMSO was added to 10mg of *Ophioglossum* in a 1.5ml eppendorf tube. The eppendorf tube was rotated overnight at a temperature of 30 – 37 °C. Appropriate amounts of the supernatant were added to culture media.

### Preparation of tripentadecanoin

Tripentadecanoin (C_48_H_92_O_6_) was obtained from Sigma-Aldrich (T4257). 5 mg of the powder was resuspended in 0.6 ml of pre-warmed (35°C) 100% ethanol and vortexed for 5 min at room temperature. The stock solution was sonicated in a water bath at 35°C for 30 min and appropriate amounts were added to culture media.

### Yeast strains

The strain used for RNAseq is s288c BY4741 wild type (yFC01: *MAT***a**, *his3*Δ1, *leu2*Δ0, *ura3*Δ0, *met15*Δ0, *ADE2, TRP1*). Strains to obtain old cells were derived from the Mother Enrichment Program (*20*) strain expressing Hsp104-GFP from its endogenous locus with deletions and *GPD_prom_-GFP-YHB1* strains were obtained according to (*10*) and are listed in Supplemental Table 2.

### Obtention of old yeast mother cells

Exponentially growing cells were diluted to OD_600nm_ of 0.02 in 25 ml SC-Full containing 1μM beta-estradiol (Sigma-Aldrich E8875) and tripentadecanoin or *Ophioglossum* whole extract, or 75 μl ethanol as a control. The yeast cultures were incubated in a shaking incubator at 30°C for 18-22 hours. Cells were pelleted (600g, 2 minutes) and resuspended in 1 ml SC-Full and supplemented with 10 μl Fluorescent Brightener 28 (Sigma-Aldrich F3543). Cells were incubated for 5 minutes in the dark and pelleted (600g, 1 minute), washed twice in 1ml SC-Full and finally resuspended in 500 μl SC-Full. 10 μl of the cells were placed on a SC-Full agar pad and imaged with a DeltaVision Elite microscope equipped with a sCMOS camera. All images were deconvolved using SoftWorx (GE Healthcare) and image analysis was performed using FIJI(*21*).

### Quantification of Yhb1-GFP and GFP-Yhb1

Exponentially growing cells were diluted in SC-Full ±tripentadecanoin (30μM) and grown at 30°C for 5 hours before imaging with a DeltaVision Elite microscope equipped with a sCMOS camera. Z-stacks were sum-projected and background was subtracted. Mean fluorescence intensity in the whole was measured. Note that GFP-Yhb1 is under a strong promoter, explaining the fluorescence level difference with Yhb1-GFP.

### Quantification of Lsm1-GFP foci in old cells

Z-stacks were max-projected and background was subtracted. All images were similarly thresholded and foci were counted in each old cell.

### RNAseq

Exponentially growing cells were diluted to OD_600nm_ 0.2 and grown for 5 hours ±tripentadecanoin (30μM) in triplicate. Cells were pelleted and plunged in liquid nitrogen. RNA extraction, library preparation and Illumina HiSeq were performed by Genewiz^®^. Sequence reads were trimmed to remove possible adapter sequences and nucleotides with poor quality using Trimmomatic v.0.36. The trimmed reads were mapped to the *Saccharomyces cerevisiae* S288c reference genome available on ENSEMBL using the STAR aligner v.2.5.2b. The STAR aligner is a splice aligner that detects splice junctions and incorporates them to help align the entire read sequences. BAM files were generated as a result of this step. Unique gene hit counts were calculated by using featureCounts from the Subread package v.1.5.2. Only unique reads that fell within exon regions were counted. The heat map from Figure 2D was generated with ClustVis (*22*). Raw counts are presented in Supplemental Table 3.

### Yeast Replicative aging

Lifespan analysis was assayed as described by Moreno et al. (*23*). Briefly, wild type MEP strain was cultured overnight in SC-Full and diluted to OD600 of 0.2 in 25ml SC-Full containing 1μM β-estradiol. Tripentadecanoin (30μM) was then added or omitted and incubated for 5hrs at 30°C, 200rpm. The culture was then diluted to OD600 of 0.01 and 500μl of this dilution was plated on YPD containing 1μM β-estradiol. Plates were incubated at 30°C for 4 days. Microcolonies were imaged using a Nikon Eclipse 50i microscope with a 10X/0.25 Nikon plan dry objective. Areas of the microcolonies were determined as in Moreno et al. (*23*) using FIJI. Data were normalised to the median of the untreated condition.

### Amyloid oligomers

Aβ_1-42_ was obtained from Bachem (ref H1368). PrP_118-135_ was obtained from Bachem (ref H-4206). Human wild-type recombinant α-synuclein was obtained from r-Peptide (ref 0101008603). Human wild-type recombinant tau (2N4R) protein was obtained from Evotec. Amylin was obtained from Bachem (ref H-7905.1000).

### Mouse primary cortex neurons

These experiments were performed by SynAging SAS on behalf of SunRegen Heathcare AG.

#### Cell culture

Cortical neurons from embryonic day 16-17 were prepared from C57Bl6/J mouse fetuses, as previously described (*24*). Dissociated cortical cells were plated (50.000 cells/well) in 48-well plates pre-coated with 1.5 μg/mL polyornithine (Sigma). Cells were cultured in a chemically defined Dulbecco’s modified eagle’s/F12 medium free of serum and supplemented with hormones, proteins and salts. Cultures were kept at 35°C in a humidified 6% CO2 atmosphere.

#### *Challenging cells with amyloid oligomers and MTT assay on cortical neurons pre-incubated with* Ophioglossum *extract or tripentadecanoin*

Before addition of vehicle or an amyloid oligomer, neurons were pre-incubated at DIV 4 with various concentrations of *Ophioglossum* extract or Tripentadecanoin for 48 h in fatty acid free medium. At DIV 6, medium was removed and cells were incubated for 24h with vehicle or 1,0 μM AβO in a final medium volume of 120 μL per well. For positive control, cells were pre-incubated with 0,05 μM DHA-ethyl ester (Sigma, D2410) for 48h before vehicle or an amyloid oligomer treatment. Cells were incubated for 24h before monitoring cell viability using the MTT assay: cells were incubated at 35°C for 1 h with MTT (Sigma, Cat #M2128-10G). For that purpose, 14 μL of 5 mg/mL MTT (solubilized in PBS) were added to each well. After incubation, medium was removed, and cells were lyzed with 150 μL DMSO for 10 minutes and protected from light. After complete solubilization of formazan, absorbance at 570 nm was recorded using a Spectrophotometer BMG Labtech Fluostar Omega. All treatments were done in triplicate.

#### *MTT assay on cortical neurons incubated with* Ophioglossum *extract or Tripentadecanoin at the same time or after cells were challenged with amyloid oligomers*

Neurons were incubated in fatty acid free medium with vehicle or an amyloid oligomer in the absence or presence of increasing concentrations of Tripentadecanoin added concomitantly to the amyloid oligomer, 3 h or 6 h after. Cells were incubated for 24h in a final volume of 140 μL per well. For positive control, cells were treated similarly in the presence of 0.1 μM HNG (S14G variant of humanin peptide). Cell viability was monitored using the MTT assay. Cells were incubated at 35°C for 1 h with MTT (Sigma, Cat #M2128-10G, Lot # MKBH7489V). For that purpose, 14 μL of 5mg/mL MTT (solubilized in PBS) were added to each well. After incubation, medium was removed and cells were lyzed with 150 μL DMSO for 10 minutes and protected from light. After complete solubilization of formazan, absorbance at 570 nm was recorded using a Spectrophotometer BMG Labtech Fluostar Omega. All treatments were done in triplicates.

#### RT-qPCR

Total RNA samples were extracted from lysates of mouse primary cortex neurons. cDNA were obtained with the Transcriptor Reverse Transcriptase kit from Roche. PCR were performed using the LightCycler system (Roche Molecular System Inc.) according to the supplier’s instructions. Transcripts analysis was done in triplicate using the primers CCCTATCTATGTGTGTCTG (forward) and TGAGGACCAAGGTATAGA (reverse) and the probe ATCTGCCTGTTGTAGTCTTAGCCTC for Neuroglobin. Data were normalized using the Rps28 gene as a control.

### Human induced pluripotent stem cells

These experiments were performed by SynAging SAS on behalf of SunRegen Heathcare AG.

Cells (HIP-Neuronal progenitors, GlobalStem, Cat#GSC-4312) were plated in 96-well plates at a density of 60.000 cells per well and culture according to supplier’s recommendations. Before experiments, cells were matured for five weeks and kept at 37°C in a humidified 5% CO2 atmosphere. Cells were incubated with vehicle or 1 μM AβO in the absence or presence of increasing concentrations of tripentadecanoin added concomitantly to AβO (T0), 3 h after AβO (T3), or 6 h after AβO (T6). Cells were incubated for 24h in a final volume of 100 μL per well. For positive control, cells were treated similarly in the presence of 0.1 μM HNG (S14G variant of humanin peptide). Neuronal loss was monitored using the detection of neuronal specific enolase by ELISA assay according to supplier’s recommendations (CloneCloud, Cat#SEA537Hu). A total of three data points per experimental condition were generated.

### NMU

These experiments were performed by IRIS PHARMA and Prof. Heping Xu at School of Medicine, Dentistry & Biomedical Science, Queen’s University Belfast, 97 Lisburn Road, Whitla Medical Building BT9 7BL Belfast, United Kingdom on behalf of SunRegen Healthcare AG. The experimental phase performed at the animal facility of Queen’s University Belfast (UK) was conducted in accordance with the ARVO Statement for the Use of Animals in Ophthalmic and Vision Research, and the study was approved by the local Animal Welfare Ethical Review Body (AWERB). NMU was obtained from Fluorochem (90%: 10% stabiliser (Acetic acid). Batch number: FCB013586). Animals were housed with 1-5 mice in each cage. All animals were maintained under a 12-hour light and dark-controlled cycle. Temperature and relative humidity were maintained at 22 ± 2°C and 60 ± 10%, respectively. Throughout the study, animals had free access to food and water. The mice were anaesthetised via intra peritoneal injection of ketamine hydrochloride (60 mg/kg, Vetoquinol UK Ltd, Northamptonshire) and xylazine hydrochloride (5 mg/kg, Bayer HealthCare, KVP pharma). The NMU solution in acetic acid (source of acetic acid) was diluted in nuclease free water (source of the nuclease-free water) to obtain a solution at 6.25 mg/mL (0.008% acetic acid) just before use. Mice received one intraperitoneal injection of NMU at a dose of 50 mg/kg (8 mL/Kg). Eighteen 8-12-week-old female C57BL/6J mice were randomized into three groups: 1) NMU + vehicle; 2) NMU + tripentadecanoin 20 mg/kg; 3) NMU + tripentadecanoin 50 mg/kg. NMU at a dose of 50 mg/kg was injected intra peritoneally to all mice; tripentadecanoin (or vehicle) was administered daily via oral gavage starting 3 days before NMU and continuing until 7 days after NMU challenge. Animals were euthanized by inhalation of CO2. After euthanasia, both eye from each animal were collected, fixed in Davidson’s solution (0.08% paraformaldehyde PFA) over-night at room temperature, rinsed in 70% ethanol for 3 hrs at room temperature, and stored at 5°±3°C. Both eyes from each animal were embedded in paraffin for histological analysis. Paraffin sections (5 to 7 μm thick) were performed along the vertical meridian and stained with hematoxylin/eosin stain. The vertical meridian included the optic nerve. Three sections per eye were examined under a standard microscope (Leica). outer nuclear layer thickness was measured every 500 μm (four points) from the optic nerve to the peripheral retina in each region of the retina (superior and inferior) using a standard microscope (Leica) operated by a single observer masked to treatment. The thickness of the outer nuclear layer was measured at each point, and the number of rows of photoreceptor nuclei was quantified. Results were expressed as the outer nuclear layer (ONL) area under the curve (AUC_-1.75 to +1.75 μm_) and number of rows of photoreceptor nuclei.

### Rhesus monkeys

Two male monkeys (*Macaca mulatta)* (monkey 9071, 5 years old, 6.6 kg; monkey 9089, 4 years old, 4.5 kg) had a family history of optic/retinal diseases and were selected into this study. Both animals did not show increased cup/disc ratio, nor increased intraocular pressure, nor increased blood glucose. They did not show glaucoma type local retinal nerve fiber layer thinness, but a general thinness in all regions of retinal nerve fiber layer. Both animals had unilateral optic atrophy localized to the right eye. Rhesus monkeys were treated with orally administered SBC003 at a dose of 5 mg/kg/day for 2 weeks, 15 mg/kg/day for 4 weeks, 30 mg/kg/day for 3 weeks, 2 weeks wash out period, 50 mg/kg/day for 4 weeks. The following parameters were evaluated during tripentadecanoin treatment: body weight, food consumption, clinical observations, clinical biochemistry and hematology, as well as ocular anterior and posterior examinations including assessment of interocular pressure, fundoscopy and ocular coherence tomography (OCT).

OCT in optic disc and macular regions: frequency: Month-4, D-8 and on D14, D28, D56, D77 and D105. Animals were anesthetized with 1:1 Ketamine:Xylazine mix (6 mg/kg ketamine, intramuscular injection), 2 drops of Tropicamide Phenylephrine Eye Drop were applied to each eye after anesthesia for pupil dilation. OCT image acquisition protocol: subject’s forehead was kept leaned against the forehead support. Following image acquisition, RNFL thickness was measured after determining the distance between the anterior and posterior surface of RNFL. The anterior and posterior surface of RNFL was detected automatically by the built-in software of Heidelberg OCT. Locating the posterior surface of RNFL manually was necessary and performed by two masked and experienced examiners. Once the anterior and posterior surface of RNFL was determined, built-in software of Heidelberg OCT showed the RNFL thickness along the circumference around the optic disc (RNFL thickness in optic disc-centered images) or the RNFL thickness in macular regions. The comparative data in the unaffected eyes served as an internal control to verify the reproducibility of OCT measurement in this study.

General Observations: During the study, the two monkeys were dosed orally. No.9071 cooperated well during the entire study; No.9089 refused to take apples from D99 and capsules were orally administered from D101. There was no major change in food intake or body weight in both monkeys. During the study, no drug-related abnormalities were observed in animals. The food intake was within the variation of food intake at this age. At the beginning of capsule dosing at week 15, No.9089 food intake of week 15 decreased to 117 g possibly due to oral dosing change from apple food-admix to capsule form, and then he fully recovered at week 16. No change was observed in biochemistry of these monkeys. Levels of ALT, AST, and BUN of monkey 9089 were slightly higher than normal limits before administration. Levels of LDL-c, ALT, and ALP of monkey 9071 were slightly higher than normal limits before administration. There was no significant change in biochemistry parameters during the study. No change was observed of the haematology of monkeys. There was no significant change in CBC parameters during the study and CBC indicators were within reference range. A pharmacokinetic (PK) study was performed on pre-dosing (8 days prior to tripentadecanoin dosing) and on Days 78, 105 at 2 hours after tripentadecanoin dosing. For monkey 9071, single point PK results showed that the exposure of tripentadecanoin increased with the dose level on D78 and D105. Tripentadecanoin plasma concentrations were tested at 6.3 ng/mL at baseline; 42.1 ng/mL at two hours after dosing on Day 78. After Day 78, the plasma concentration fluctuated between 47.6-86.4 ng/mL, showing a trend of increased exposure with an increased dose. For monkey 9089, the plasma concentration of tripentadecanoin was not increased with the increased dose. It was reduced to 27.1 ng/mL or close to baseline regardless of the increased dose.

All procedures in this protocol followed the Animal Welfare Act, the Guide for the Care and Use of Laboratory Animals, and the Office of Laboratory Animal Welfare, and approved by IACUC of Sichuan Primed Shines Bio-tech Co., Ltd.

